# Temporal masking and rollover in the neural code for speech with and without hearing loss

**DOI:** 10.1101/2022.11.10.515823

**Authors:** Chengjie G. Huang, Shievanie Sabesan, Nicholas A. Lesica

## Abstract

Natural sounds, such as speech, are complex time-varying waveforms containing information critical to how we communicate with each other and navigate the external world. Hearing loss results in a breakdown of this information and causes distortions in the neural code. As a result, perception of complex sounds such as speech is compromised. This problem is further complicated by the fact that sound intensity varies in natural settings, both in quiet and in noisy backgrounds. Somewhat paradoxically, despite increased audibility at high sound intensities, perception and discrimination of speech is actually diminished, especially in the presence of background noise. This phenomenon is known as rollover of speech and its neural basis is poorly understood in both normal-hearing listeners and hearing-impaired listeners. Here we performed in-vivo electrophysiology in awake and anaesthetized Mongolian gerbils *(Meriones Unguiculatus)* to investigate how hearing loss affects the neural encoding of speech. We presented 22 Vowel-Consonant-Vowel (VCV) syllables to the gerbil and recorded neural responses from the inferior colliculus (IC). We used a k-nearest neighbor neural classifier to investigate whether IC neurons could discriminate between different consonants in normal hearing (NH) and noise-exposed hearing-loss (HL) animals. We found that neural correlates of perceptual rollover were present in the IC and that performance in discrimination decreased when VCVs were presented in background noise when compared to in quiet. Furthermore, we found that forward masking played a prominent role in shaping neural responses and discrimination between various consonants in NH and HL animals. These results suggest there is a critical trade-off in listening between audibility and rollover mediated by temporal masking.

## Introduction

Natural sound processing used in everyday situations is fundamental to how humans interact with each other and with the external world. Natural sounds, such as speech, are complex time-varying waveforms containing rich spectrotemporal information critical to communication. Notably, the lack of access to this critical information in sensory disorders, such as in sensorineural hearing loss (SNHL) decreases the quality of life for those affected and has become a global socioeconomic burden. In general, SNHL is thought to be caused by a breakdown in the neural signals carrying sound information, or neural code, leading to distorted perception of sounds ((Zhong et al., 2014; Henry et al., 2016; Lesica, 2018). Within this context, there are many understudied phenomena which are thought to have distinct neural mechanisms that are poorly understood to date. One such phenomena is the observation that listening to speech in noise with increased sound intensity, and by extension supposed increased audibility, paradoxically leads to decreased sound perception. This is known as rollover of speech and it has been observed even in normal-hearing individuals (Hornsby et al., 2005; Summers and Cord, 2007). However, the study of this phenomenon has been largely limited to psychophysical studies in humans and the underlying central neural mechanisms that lead to rollover are unknown in both normal-hearing and hearing-impaired individuals.

Hearing impaired individuals such as those with SNHL have poor speech-in-noise perception even at high sound intensities. Therefore, it is critical to investigate the underlying neural mechanisms which lead to rollover in order to gain a clear understanding of how increased intensity impairs perception and how the neural code is distorted at higher sound intensities in both normal-hearing and SNHL individuals. The neural code carries essential information through successive stages of the auditory system from the auditory nerve to the auditory cortex. At the level of the periphery, complex distortions caused by high intensities and/or SNHL may not yet manifest, while higher brain areas such as the thalamus and cortex, involve contextual and behavioral factors which complicate studying the distortions in the neural code due to modulations from cognitive functions (Fritzsch et al., 2015; Jayakody et al., 2018). Here, we chose to study rollover in the inferior colliculus (IC), which is the midbrain hub of the central auditory pathway and acts as the essential relay between early brainstem and the thalamus. The neural activity present in the IC reflects the integrated effects of processing of several peripheral pathways while the neural response is still primarily determined by the acoustic features of incoming sounds ((Moore et al., 1983; McAlpine et al., 1997).

We performed large-scale electrophysiology recordings Mongolian Gerbils (*Meriones Unguiculatus*). The gerbil is an attractive animal model to investigate age-related hearing loss because of its well-characterized physiology and anatomy (Henry et al., 1980; Mills et al., 1990) and its similar sensitivity to humans for frequencies between 1 and 4 kHz (Ryan, 1976). We performed IC neural recordings in both normal hearing and noise-exposed hearing loss (HL) animals to assess the neural effects of rollover. We presented a set of 22 Vowel-Consonant-Vowel (ae-Consonant-ae) syllables across a spectrum of increasing sound intensities both in quiet and various background noise levels and used a neural classifier to simulate perception. We hypothesized that the neural response to the consonants would vary between normal hearing and HL animals at different sound intensities and would also change across different speech-to-noise ratios (SNR). In addition, we predicted that IC neurons would exhibit rollover as decreased neural classifier performance when the speech is presented in noise and that this effect would be more prominent in normal hearing animals than in HL animals. Finally, we hypothesized that rollover of speech at high sound intensities arises from temporal masking effects between the vowel-consonant and consonant-vowel transition. We predicted that isolating the individual components of the syllable would improve neural classifier performance, demonstrating a mitigation of the effect of rollover.

## Materials and Methods

### Animals

Experiments were performed on 9 young-adult gerbils of both sexes that were born and raised under standard laboratory conditions. Four of the gerbils were exposed to noise when they were 10–12 weeks old. Experimental recordings were performed for all of the gerbils when they were 14–18 weeks old. All protocols performed were approved by the Home Office of the United Kingdom under license number 7007573. All experimental control and data analyses were performed using custom code in MATLAB R2018a (Mathworks).

### Noise Exposure

Mild-to-moderate sensorineural hearing loss was induced by exposing anaesthetized gerbils to high-pass-filtered noise with a 3dB per octave roll-off below 2 kHz at 118dB SPL for 3h (Suberman et al., 2011). For anesthesia, an initial injection of 0.2ml per 100 g body weight was given with fentanyl (0.05mgml^-1^), medetomidine (1mgml^-1^) and midazolam (5mgml^-1^) at a ratio of 4:1:10, respectively. A supplemental injection of approximately 1/3 of the initial dose was given after 90min. The internal temperature was monitored and maintained at 38.7 °C. Successful sensorineural hearing loss was confirmed with auditory brainstem responses (ABR) performed on the day of electrophysiological recordings.

### Electrophysiology

Animals were placed into a sound-attenuated chamber and anaesthetized for surgery with an initial injection of 0.65ml per 100 g body weight of ketamine (100mgml^−1^), xylazine (20mgml^−1^) and saline in a ratio of 5:1:19, respectively. The same solution was infused continuously during recording at a rate of approximately 2.2μlmin^−1^. The internal temperature of the animal was monitored continuously by a temperature probe and maintained at 38.7 °C. A small metal rod was mounted onto the skull and used to secure the head of the gerbil in a stereotaxic device. Incisions were made in order to reveal the skull, where a small metal rod was attached anterior to bregma and secured to a stereotaxic device. After aligning the skull, two craniotomies were made lateral to the midline and rostral to lambda over the recording site where the inferior colliculus (IC) was accessible. Incisions were made in the dura mater, and a 256-channel multi-electrode array (NeuroNexus) was inserted into the central nucleus of the IC in each hemisphere. The arrays were custom-designed to maximize coverage of the portion of the gerbil IC that is sensitive to the frequencies that are present in speech. As the arrays approached the central nucleus of the IC, they were moved slowly into the optimal location for recording based on an online analysis of the site FRAs to observe a range of BFs relevant to speech syllables.

### Auditory Brainstem Response

The protocol for the ABRs performed were based on Poloneko and Maddox (2019) and adapted for gerbils. ABRs were performed prior to every electrophysiology experiment to ensure sensorineural hearing loss had occurred in the animal from the noise exposure protocol. Once the animals were under anesthesia for recording, subdermal needles were used as electrodes with the active electrodes placed behind the ear over the bulla (one on each side), the reference placed over the nose, and the ground placed in a rear leg. Recordings were bandpass filtered between 300 and 3000 Hz. Randomly timed tones were presented at multiple frequencies simultaneously and independently to each ear. The tone frequencies were 500, 1000, 2000, 4000, and 8000 Hz. Each tone was 5 cycles long and multiplied by a Blackman window of the same duration. Tones were presented at a rate of 40 per s per frequency with alternating polarity for 100 s at each intensity. The activity recorded in the 30 ms following each tone was extracted and thresholds for each frequency were defined as the lowest intensity at which the root mean square (RMS) of the median response across presentations was more than twice the RMS of the median activity recorded in the absence of sound.

### Sounds

All sounds were delivered to speakers (Etymotic ER-2) coupled to tubes inserted into both ear canals along with microphones (Etymotic ER-10B+) for calibration. The sounds were amplified via a power studio amplifier (RA150, Alesis). The full set of sounds played is described in detail below:

1. Rate Level Function: 50 ms broadband noise bursts at intensities ranging from 1 dB SPL to 85 dB SPL in 6 dB steps with a 175 ms pause between sounds. Sounds were presented 32 times each in a random order. This sound was played approximately every hour to ensure that the intensity sensitivity threshold remained stable.
2. FRA Tone set: 50 ms tones with frequencies ranging from 300 Hz to 10000 Hz in 0.25-octave steps and intensities ranging from 4 dB SPL to 85 dB SPL in 9 dB steps with 5ms cosine on and off ramps and a 175 ms pause between tones. Tones were presented 8 times each in a random order.
3. PST Tone set: 50 ms tones with frequencies ranging from 300 Hz to 10000 Hz in 0.5-octave steps at either 60 or 85 dB SPL with 5 ms cosine on and off ramps and a 175 ms pause between tones. Tones were presented 128 times each in a random order.
4. Vowel-Consonant-Vowel (VCV) Syllables: Speech syllables in the configuration of ‘aeXae’ from a single adult female talker were presented in 22 variations corresponding to 22 different consonants and were each repeated 32 times across each sound intensity. The 22 consonants consisted of a variety of consonants in the English language including */p/, /t/, /k/, /f/, /s/, /b/, /d/, /g/, /v/, /z/, /m/, /n/, /w/, /r/, /j/, /l/, /θ/, /* ð */,/ dʒ/, / t⎰/, / ʒ/, / ⎰/*. Each token of VCV syllable were presented at sound intensities of 46, 54, 62, 70, 78, 86, and 94 dB SPL to allow a full range of investigation into sound level dependent neural responses. We additionally presented these sounds in quiet and in at various SNRs ranging from -2 dB to +10 dB SNR with background noise described below in 6). All trials were considered for both quantification of neural responses as well as neural classification for training and testing.
5. Isolated Consonants: Using the VCV stimuli waveforms above from, two cutoff points were made using spectral analysis to indicate the boundaries between Vowel 1, the Consonant, and Vowel 2. The waveform between the two cutoff points were extracted to use as the isolated “gapped” stimuli which were additionally presented to the animal on separate blocks. Each token of the isolated consonant was presented at the same sound intensities as 4) as well as in quiet and in background noise described in 6) below.
6. Background Noise: Continuous speech from 16 different British English speakers from the UCL Scribe database (https://www.phon.ucl.ac.uk/resource/scribe) was summed to create speech babble. The intensity of the babble was set based on the intensity of the VCV syllables to achieve the desired speech-to-noise ratio.

## Data analysis

### Representation of Neural Responses

Multi-unit activity (MUA) was measured from recordings from each of the 512 sites on the electrodes. The raw trace was high-pass filtered with a cutoff frequency of 500 Hz, followed by taking the absolute value, and subsequently low-pass filtered with a cutoff frequency of 300 Hz. A thresholding process was then applied to produce discrete pseudo-spiking time-varying signal segregated into each time bin at the downsampled sampling frequency of 625 Hz. The resulting MUA was used for the classification analyses purposes described below.

### Classification of Neural Responses

Single trials were classified using the k-nearest neighbors algorithm with *k* = 1 as follows:

1) Multi-units were grouped in populations by selecting randomly across the pool of all recording sites across all animals in each group (i.e. Control or HL) in sets of 50. 2) The time window to be analyzed occurred 72 ms from the first cutoff time indicating the beginning of the consonant in each trial. 3) The population response to a single trial of each VCV syllable was removed from the full set of all responses. 4) The Euclidean distance between the removed responses and each of the remaining responses in the set was computed. 5) Each removed response was classified as having been elicited by the same consonant as the response in the remaining set to which it was closest. Ties were broken by selecting the template response with the lower median Euclidian distance across iterations of the bootstrapped trials. Confidence intervals were estimated by bootstrapping the with replacement during the classification 100 times.

### Decoding of “Stitched” Responses

In addition to classifying using the ungapped neural responses and the gapped isolated consonant neural responses, we performed classification using responses stitched from the two types of stimuli. Specifically, we extracted the response to the isolated consonant in its entirety and replaced the response to the ungapped consonant using the specific cutoff times for each consonant. We then performed our k-nearest neighbors classification on the same time window of 72 ms. All methodologies in the decoding process occurred in the same process as mentioned above.

### Decoding of “Hybrid” Responses

To investigate whether the temporal masking effect was either forwards or backwards masking, we additionally created “hybrid” consonants using the first 40 ms of the isolated consonant neural response and replacing the response in that time window from the beginning of the consonant neural response in the ungapped condition. These consonants would therefore remove the influence of forward masking, but retain much of the original response in the ungapped condition with the influence of backward masking intact. We then performed our k-nearest neighbors classification on the same time window of 72 ms.

## Results

To study the effect of rollover of speech at high sound intensities, we performed large-scale electrophysiology recordings from populations of neurons in the IC with custom-designed electrodes with a total of 512 channels spanning both brain hemispheres in gerbils (Fig. 1A). These recordings resulted in small subsets of neurons recorded on each channel of the electrode, or multi-unit activity (MUA), which we collectively analyzed in detail to study as neural responses to auditory stimuli presented (Fig. 1B). To induce sloping mild-to-moderate sensorineural hearing loss, we exposed young-adult gerbils to broadband noise (118 dB sound pressure level (SPL) for 3h). We confirmed elevated hearing thresholds to pure tones by performing parallel auditory brainstem response (ABR) recordings (Polonenko and Maddox, 2019) one month after noise exposure as well as on the day of the electrophysiology recording to confirm the results (Fig. 1C). Compared with normal hearing animals, the noise exposed (HL) animals demonstrated elevated thresholds of 20–30 dB at low frequencies (∼1 kHz) to 40–50 dB at high frequencies (∼8 kHz). This was confirmed by both on-line and offline analysis of frequency responses area curves (FRAs) obtained from the MUA data (Fig. 1D), which tested the responses of populations of neurons at each electrode channel to systematically presented pure tone frequencies and intensities.

**Figure 1.**
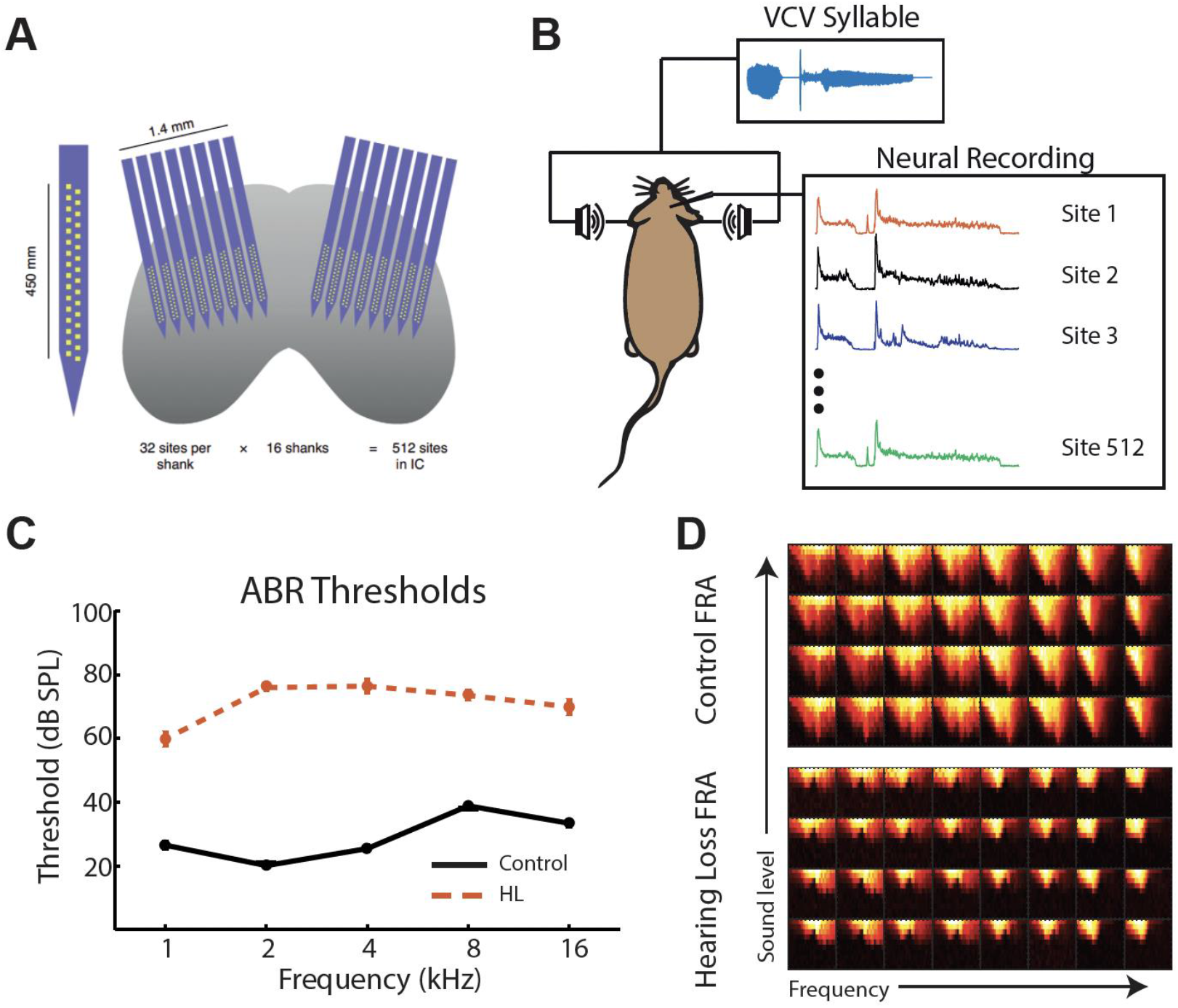
Neural and ABR recordings in Mongolian gerbils. A) Schematic of neural electrophysiology recordings made in the IC of gerbils with two custom 256-channel electrodes. B) Schematic of auditory stimulus (blue) delivery via tube-coupled speakers and neural responses recorded across each of the 512 sites on the electrode (colored traces). C) ABR thresholds of control (black) and HL (orange) animals. D) Frequency-response area plots of control (top) and HL (bottom) animals. Each box corresponds to one recording site on the electrode with the x-axis indicating increasing frequency from left to right and y-axis indicating increasing sound level from bottom to top.

To study the neural mechanisms which underlie rollover of speech, we presented speech stimuli in the form of Vowel-Consonant-Vowel (VCV) syllables with a single vowel ‘ae’ combined with 22 different consonants: */p/, /t/, /k/, /f/, /s/, /b/, /d/, /g/, /v/, /z/, /m/, /n/, /w/, /r/, /j/, /l/, /θ/ (xt), / ð / (xd),/ dʒ/ (xj), / t⎰/ (xc), / ʒ/ (xz), / ⎰/ (xs)*, all spoken by a single female speaker. The 22 consonants corresponded to a range of sounds in the English language and was used in Hornsby et al., (2005) to test rollover of speech in human participants. In addition, these nonsense syllables are commonly used in other psychophysics studies in humans to assess perception and discrimination abilities in hearing (Repp, 1977; Klaassendon and Pols, 1983; Dubno et al., 2012). We presented 16 trials of each of the 22 syllables shuffled in a random order, summing to a total of 352 tokens in each block. We then presented each block at different sound intensities ranging from 62 – 94 dB (see Materials and Methods for details) to assess how the neural response to each consonant syllable is differentially encoded at increasing sound levels and the corresponding effects of rollover.

### IC Neurons respond robustly to speech syllable sound stimuli

While presenting “ae” VCV syllables to the gerbil at various intensities, we recorded neural responses in the IC. We extracted the MUA by first filtering the raw response (see Materials and Methods) and subsequently applying a threshold to be used for offline processing. The thresholding process produced a discrete pseudo-spiking time-varying signal segregated into each time bin at the downsampled sampling frequency of 625 Hz. To visualize the overall population activity, we averaged the thresholded MUAs across channels to produce a peri-stimulus time histogram (PSTH) for each token and trial of the stimulus. The PSTHs demonstrated robust responses to the VCV syllable stimuli, especially in quiet for both control and HL animals. The PSTH responses reliably tracked the stimulus and different consonants elicited differential PSTH responses (Fig. 2). As expected, the amplitudes of the PSTHs obtained from HL animals were lower than those in control animals, in particular at low sound intensities where HL animals had reduced audibility to the sound presented. However, in general, the response amplitudes of the PSTHs tended to increase with increasing sound intensity for both control and HL animals.

**Figure 2.**
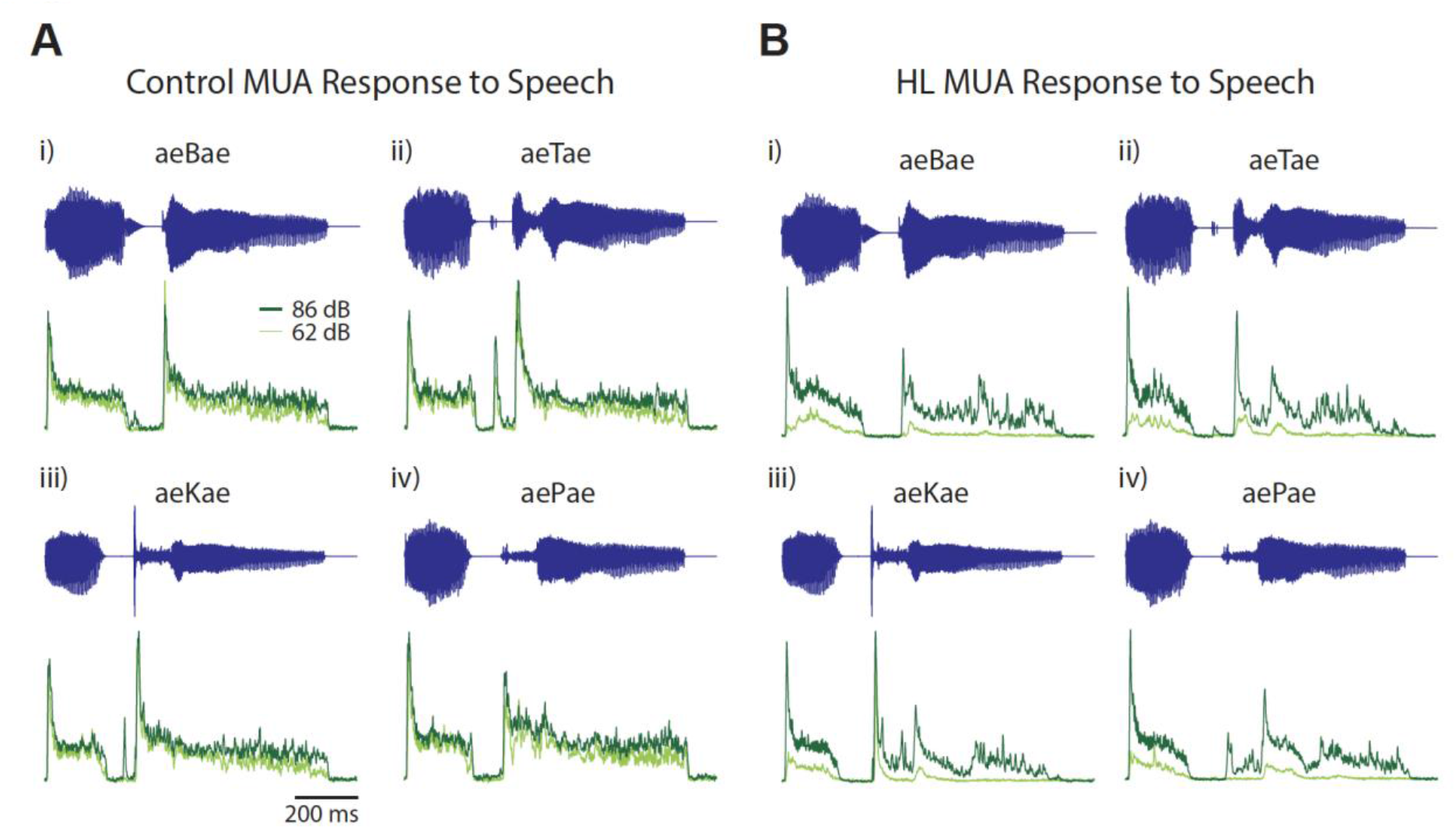
Multiunit activity (MUA) responses to speech syllables. A) Averaged MUA PSTH responses in a control animal to 4 sample vowel-consonant-vowel speech syllables (blue): i) ae/B/ae, ii) ae/T/ae, iii) ae/K/ae, and iv) ae/P/ae at 62 dB SPL (light green) and at 86 dB SPL (dark green). B) Same as A) except for a noise exposed hearing-loss (HL) animal.

Following presentation of the syllables in quiet (i.e., infinite SNR), we then presented the syllables in background babble noise at SNRs at -2, +2, +6-, and +10-dB SNR. For both control and HL animals, the PSTH responses became noisier and demonstrated lower overall response amplitudes at -2 dB SNR background noise. As SNR was increased to +10 dB SNR, there was a graded increase in the PSTH response amplitude to the elements of the syllable and reduced responses to the background noise. The responses at +10 dB SNR were not significantly different from the responses under quiet conditions (data not shown) and this trend was modulated by the increasing intensity mentioned above.

### Neurons in the IC can correctly classify consonants in quiet

While intuitively it appeared that the PSTHs obtained from increasing intensities correlated with increased amplitudes in the neural response, this may have either a positive or negative effect on discrimination between different consonants. To identify whether consonants can already be distinguished at the level of the IC, we used a neurogram-based classifier to assess how discrimination varies as a function of increasing intensity as well as background noise. Our recordings yielded ∼1500 high-quality electrode sites from each of our control and HL gerbil groups. We then randomly allocated those recording sites into groups of 50 sites which we considered one “experiment.” Using the MUA from each of the 50 sites, we constructed 2-D neurograms with each row’s x-axis representing time and the y-axis representing the MUA response over 50 sites (rows). Each neurogram represented the time window of the consonant presented in one token of speech syllables presented to the gerbil. We then iterated through each token’s neurogram, while removing all tokens associated with the trial number of that particular syllable stimulus. We used a k-nearest neighbor (KNN) algorithm to find the closest neurogram in Euclidian distance. This would yield one closest neurogram which corresponded to a syllable. If the KNN neurogram’s syllable was the same as the token neurogram in question, then this would be a considered a correct classification, and incorrect otherwise. Using this method, we generated confusion matrices over the average of the bootstrapped tokens and trials, which provided an accurate percent correct for each of the 22 consonants.

When the speech syllables were presented in quiet (Fig. 3A left), control animals yielded an overall percent correct which approached maximum performance even at the lowest sound intensity (91.0% at 46 dB), and >99% at higher intensities. In HL animals (Fig. 3A right), the overall percent correct approached maximum performance only at >= 62 dB sound intensity. This is expected given that HL animals still experience threshold shifts and audibility issues associated with hearing loss. However, once the sounds presented reach audible intensities for the HL animals, classification of consonants were similar to control animals.

**Figure 3.**
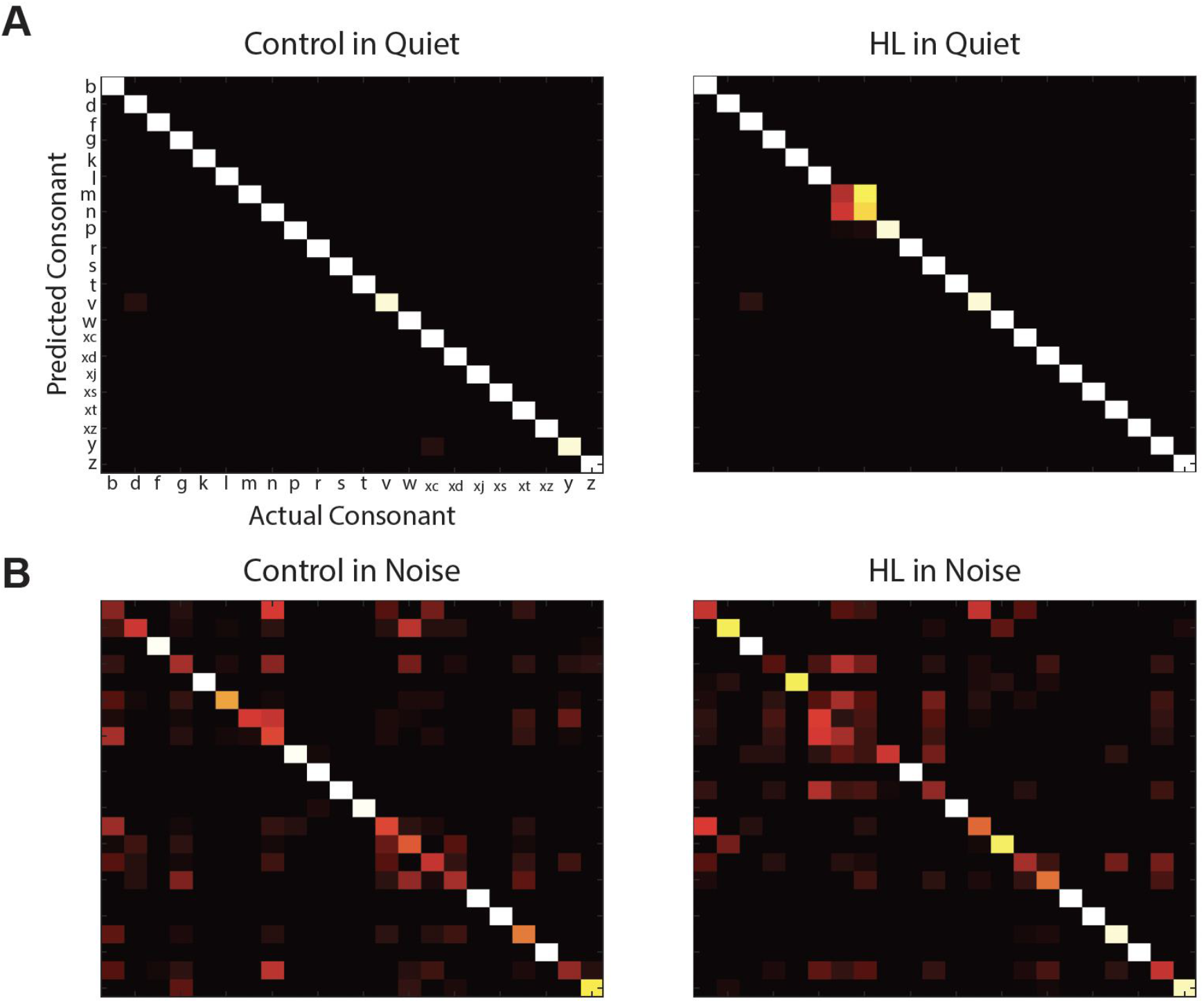
KNN classification of consonants. A) Example confusion matrices for control (left) and HL (right) animals for speech in quiet condition. Colours indicate percent correct for matches between predicted consonant and actual consonant presented. B) Same as A) except for in background noise condition at +2 dB SNR.

### Consonant classification performance is reduced in noisy conditions

Although IC neurons in both control and HL animals perform reasonably well in classifying consonants in quiet, this becomes much more complicated when the speech is presented in background noise. We presented the same speech syllables in -2, +2, +6, and +10 dB SNR while also increasing the sound intensity to assess the effect of rollover. We found that -2 dB SNR resulted in noisy neural signals which resulted in a condition which was too difficult to accurately assess classification. Furthermore, the classification performance at +6 dB and +10 dB SNR were not significantly different than the performance in quiet conditions (data not shown), and as such it appears that they were equivalent to infinite SNR demonstrating the robustness of the neural code with a sensitive range where rollover operates. For the sake of simplicity, we will therefore assess the effect of rollover with speech-in-noise designated as the results found with +2 dB SNR conditions.

The PSTHs for +2 dB SNR neural responses were strikingly different visually from the PSTHs from in quiet conditions and this varied as a function of different consonants as well as control and HL animals. We hypothesized that the neural basis of rollover of speech were evident in the IC, then the neural activity should reflect a consistent decrease in classification performance as sound intensity increased. For control animals, this was indeed the case as the classification performance increased to a peak at 62 dB and subsequently decreased in performance as sound intensity increased (Fig. 4A top). This demonstrates that already in the neural populations of the IC, the ability for neurons to distinguish between consonants becomes a problem as a result of the confusion between consonant sounds when they are presented in presence of background noise. We also hypothesized that should the classification performance decrease, then the confusion between consonants should reflect those found perceptually due to rollover of speech. While assessing the confusion matrices (Fig. 3B and Fig. 4A top), it becomes clear that the same consonants were being confused by the neural populations during classification as those found in human psychophysics perceptual studies, in particular the */l/, /m/, /n/*, and */v/, /w/, /xc/*, and */xd/* consonants. These consonants become further confused amongst each other as sound intensity is increased in the presence of background noise, giving rise to lower performance (Fig. 4B) and hence the observation of a clear neural correlate of rollover.

**Figure 4.**
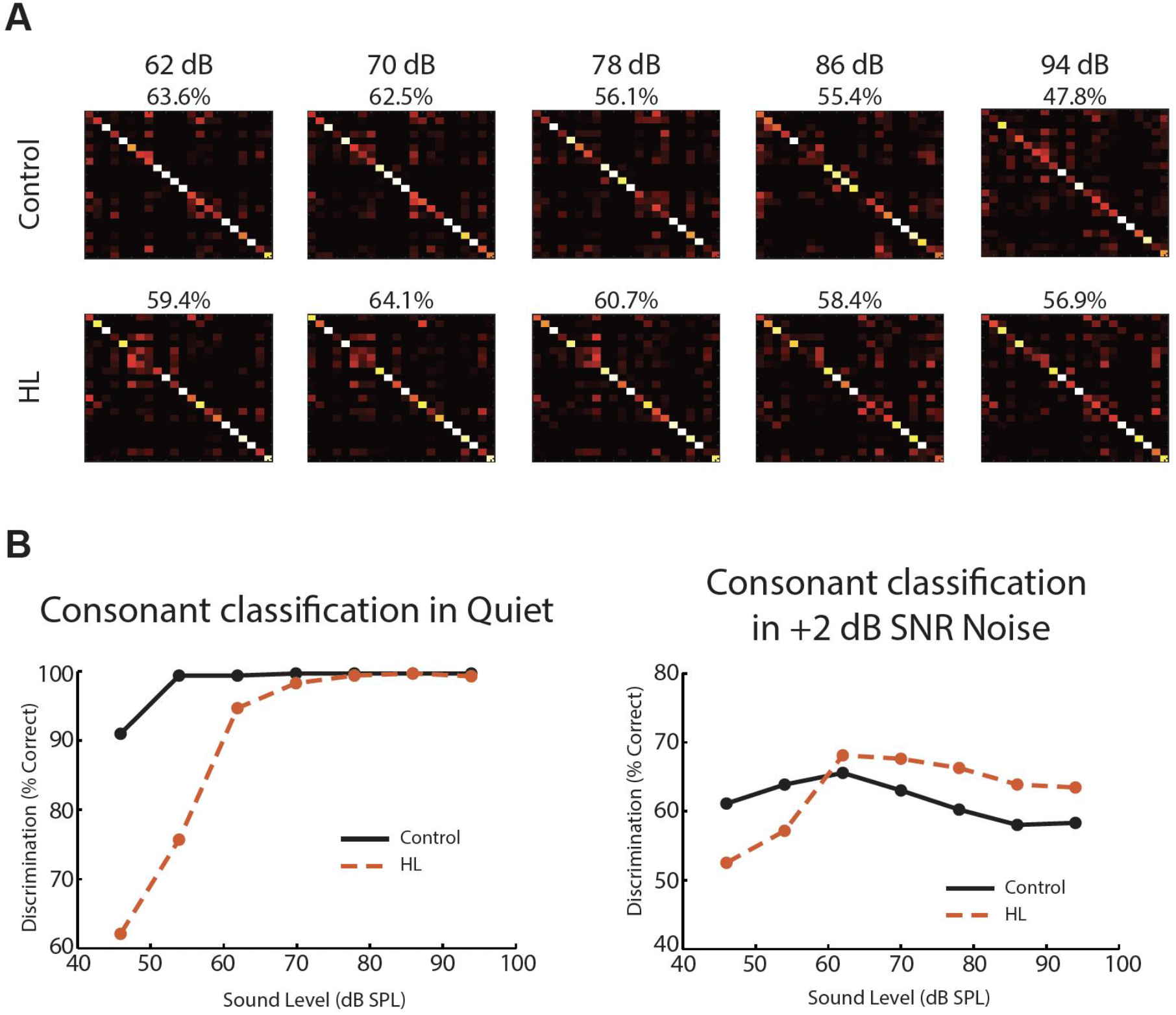
Neural correlate of rollover occurs in the IC. A) Example confusion matrices for a control (top) and HL (bottom) animal across 5 different sound levels (62-94 dB SPL). B) Consonant classification tuning curves across increasing sound levels for in quiet condition (left) and in +2 dB SNR background noise (right) for control (black) and HL (orange) animals.

Surprisingly, when the same set of syllables were presented to the HL animals in the presence of background noise, the effect of rollover was a lot less prominent in the neural responses (Fig. 4A bottom). As with the control animals, classification performance peaked at 62 dB with a lower rate of performance decrease per dB when compared to the control animals. Additionally, a surprising result was that the neural classification performance of the HL animals was actually higher than the control animals at sound intensities >62 dB (Fig. 4B right, compare solid with dashed). This will be addressed in the next section below. However, the classification performance was still overall decreased compared to when sounds were presented in quiet (compare Fig. 4B left with Fig. 4B right). Finally, as with the case in control animals, the consonants which were being confused by the neural populations during classification were similarly found (Fig. 4A bottom). In conclusion, these results suggest that HL animals were resistant to the effects of rollover and this is reflected by the key finding that the decrease in classification performance dropped more slowly as sound intensity of the stimuli was increased when compared with control animals.

### Temporal masking effects underlie the deterioration of the neural code at high sound intensities

So what is the underlying mechanism responsible for the strong effect of rollover in control animals when listening to speech in the presence of background noise in contrast to HL animals? We hypothesized that the border between consonants and vowels became a vulnerable neural coding space where the transition between the phoneme components could possibly induce masking effects. This is to say that there is the possibility of forwards masking of vowel 1 onto the consonant and backwards masking onto the consonant by vowel 2 in our *ae/C/ae* paradigm. To test this possibility, we isolated the consonants from the vowels preceding and following them with periods of silence on either side to replace the vowels. To this end, we have essentially created a consonant which is infinitely gapped from the vowel components in the syllable, removing any possible influence of forwards or backwards masking. For simplicity’s sake, we will call the original non-spliced syllable “ungapped” and the isolated consonant “gapped”.

The PSTHs for the gapped neural responses demonstrated that the consonant responses were larger in amplitude when compared to the ungapped neural responses (Fig. 5A) for both control and HL animals, which suggested there was a release from the diminished robust responses for individual components of the syllables due to the possibility of temporal masking. This effect appeared in several of the consonants in the dataset demonstrating an “unmasking” of consonant and vowel responses when the consonants were presented in isolation. We then wanted to find out whether temporal masking had a direct effect on rollover and the ability for correct neural classification of consonants.

**Figure 5.**
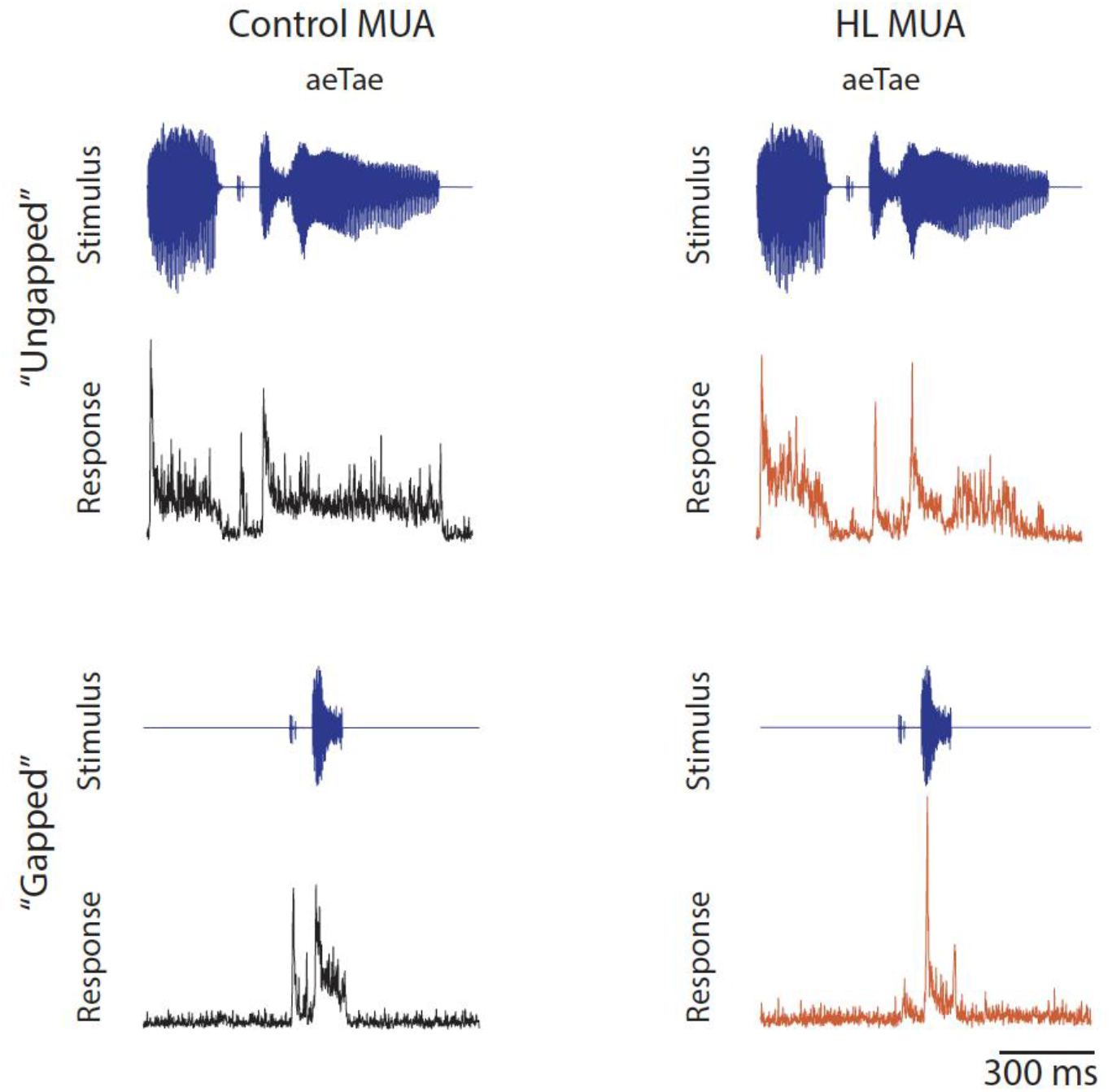
Isolated “gapped” consonants elicit neural responses without temporal masking. Example averaged MUA PSTH responses to an ungapped stimulus (top) and a gapped stimulus (bottom) for a control (black) and a HL (orange) animal.

We extracted the gapped neural response for the entire duration of each consonant and replaced the neural response in the ungapped neural response corresponding to the same consonant across the 16 trials (Fig. 6A). For simplicity’s sake, we have termed these responses “stitched” to distinguish them from the original ungapped responses. This essentially created a neural response where the vowel responses were still intact, while the consonant response was unaffected by the potential temporal masking effects of either vowel. By reclassifying in a similar manner as before, we make an equal comparison while accounting for the various durations of the consonant responses. After performing the KNN classification, we found that the performance improved overall for control animals when compared to the ungapped condition, while rollover was still present (Fig. 6B compare black curves). For HL animals, however, the difference between performance in the ungapped and stitched conditions were much smaller overall, and the removal of temporal masking seemed to mitigate the weak rollover that was present in the ungapped condition (Fig. 6B compare orange curves).

**Figure 6.**
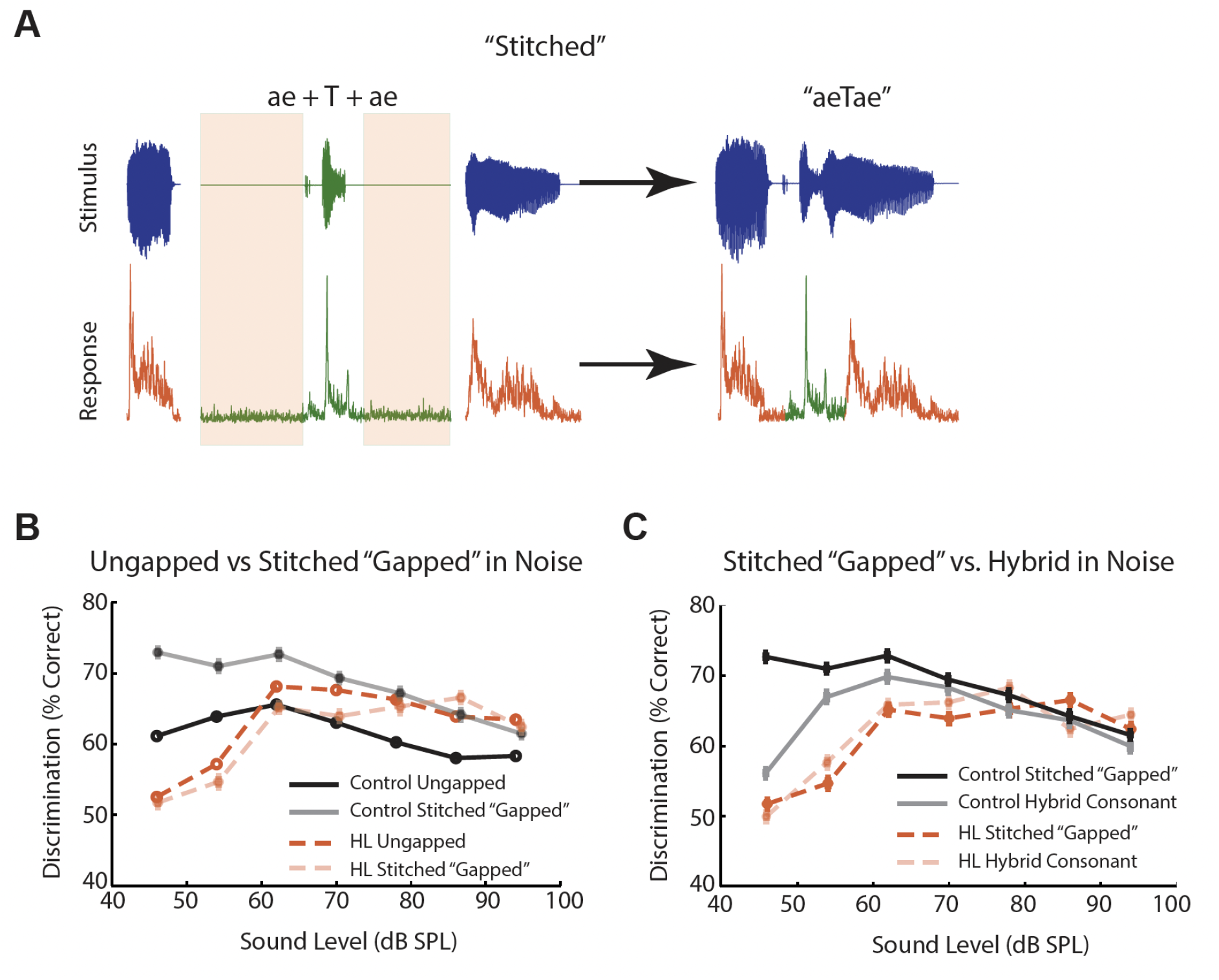
Forward masking plays a prominent role in shaping the neural correlate of rollover. A) Methodology in which “stitched” neural responses were created by extracting the neural response to the full consonant from the gapped neural response (green) and replacing its corresponding time window in the ungapped neural response with vowel responses (orange) intact. B) Comparison of averaged consonant classification tuning curves for ungapped neural responses (solid) and stitched neural responses (shaded) for control (black) and HL (orange) animals. C) Comparison of averaged consonant classification tuning curves for ungapped neural responses (solid) and hybrid consonant neural responses (shaded) for control (black) and HL (orange) animals.

Finally, we wanted to investigate whether the effects of temporal masking were caused by forward or backward masking from either vowel 1 or vowel 2 respectively. We predicted that it is unlikely that different utterances of vowels following the consonant could give rise to influences which led to the confusion between consonants, and thus predicted that it was most likely a forward masking effect. To test this, we similarly took the stitched method used above, with the key difference of only extracting the first 40 ms of the gapped neural response to replace the same corresponding time window in the ungapped neural response.

This essentially created a “hybrid” consonant where the first 40 ms of each consonant was without influence from temporal masking, while anything following would be the original ungapped response which was possibly susceptible to backwards masking from the second vowel. We predicted that if the curves did not change, then it is likely that only forward masking played a role in the mechanism underlying rollover. Indeed, after classification, we found that the performance curves were similar between the stitched responses and the hybrid consonant responses, at least for intensities above 50 dB SPL (Fig. 6C). These results above confirm our predictions that forward masking from the first vowel onto the consonant was largely responsible for the difference between performance in the ungapped and stitched conditions.

## Discussion

The primary goal of this study was to identify how rollover of speech occurs and determine if there was a neural correlate of this phenomenon. Furthermore, if rollover was already observed in the auditory pathway, was there a neural mechanism which could be exploited to mitigate the effect of rollover? Our results suggest that this was indeed the case as “neural rollover” was already observed in the IC when we analyzed the neural activity in response to speech syllables presented in noise at increasing sound intensities. In addition, when we removed the possible temporal masking effects present in the syllable between vowels and consonants and vice-versa, we found that performance for control animals was dramatically improved, while performance for HL animals remained similar overall but with decreased rollover. Thus, we not only deduced a neural correlate of the rollover phenomenon in the brain, but also demonstrated that the role of temporal masking differs across hearing conditions.

To date, the mechanisms which underlie rollover have been poorly understood and the majority of the literature have been focused on human psychophysics studies where perceptual performance was used to directly measure perception of speech-in-noise at high sound intensities. Despite this, the studies have been extremely limited in their datasets and testing groups. It was assumed that when sound intensity is increased, for sounds in general, that the temporal envelope changes as a result of compression in the basilar-membrane response played an important role in shaping the psychometric function of speech, as speech perception was primarily thought to be dependent on envelope cues (Moore and Glasberg, 1989; Glasberg and Moore, 1992). This hypothesis was directly tested in speech recognition tasks. For example, when consonant-vowel syllables and sentences were degraded using noise vocoders to introduce large changes to the speech envelope, it was found that perceptual scores of normal-hearing listeners increased significantly from 45 to 60 dB SPL and decreasing scores and feature transmission from 60 to 85 dB SPL (Dubno et al., 2012). These results made predictions that the spectral features in the speech envelope were indeed highly important. In addition, another study found that speech recognition scores showed larger and more consistent decreases at high levels (i.e., greater rollover) for high-frequency speech materials in comparison to low-frequency speech materials (Studebaker et al., 1999; Molis and Summers, 2003). This further lends to the idea that rollover is a multi-dimensional problem spanning from basilar membrane compression to the successive processing in the auditory pathway.

One key takeaway idea from these studies was that these speech recognition experiments were performed on normal-hearing listeners, who were susceptible to the effects of rollover. We found similar results in control gerbils in the present study, as presenting speech syllables in the presence of background noise resulted in decreasing neural classification performance with increasing sound intensities. Therefore, it appears that there are several spectrotemporal factors at a fundamental level which are varied with increasing sound intensity, leading to the rollover phenomenon as normal-hearing listeners are not excluded from experiencing it. This then leads into the question of whether the rollover effect is worse for hearing-impaired individuals. One would expect that should rollover be a simple spectrotemporal problem (such as low-frequency vs. high-frequency effects), that for listeners with severe SNHL at high frequencies, speech at levels required a achieve audibility at that range would often fail to improve performance at all (Hogan and Turner, 1998; Ching et al., 2001). On the contrary, the amplification of low-frequency speech to very high levels could potentially benefit hearing-impaired listeners as this range of speech materials is less susceptible to the effects of rollover based on the studies mentioned above. Interestingly, one study found that for high-frequency stimuli in the range of 75 to 87.5 dB increase resulted in a prominent rollover effect for normal-hearing listeners, but performance remained constant for hearing-impaired listeners (Summers and Cord, 2007). The noise-induced distortions at high frequencies at high sound intensities are perhaps greater for normal-hearing listeners than for hearing-impaired listeners, where hearing-impaired listeners may benefit from increased audibility while mitigating the negative consequences of rollover. In the present study, we found this similar benefit in the HL animals where classification performance remained stable with only minor decrease as sound intensity increased. These results suggested that auditory perception, and to the neural coding extent in the brain, is a balancing act between audibility and distortions (such as those found in rollover). Because HL animals, and perhaps hearing-impaired individuals experience reduced audibility, amplification of sounds to high sound intensities would not simultaneously drive large distortions, with a net benefit effect on perception. In contrast, the control animals, and normal-hearing individuals are already at a baseline level of audibility where amplification is not needed for perception at a comfortable listening level, the increase in sound intensity induces the distortions which are consequentially detrimental for perception.

Finally, one needs to address the temporal component of neural coding as speech occurs in a limited timescale for processing and recognition. The problem of rollover should therefore not be constrained to frequency of the speech material as any combination of syllables with various low-frequency (vowels) and high-frequency (consonants) can exist in any given word, phrase, or sentence. The present study posits a universal temporal mechanism which becomes problematic in conjunction with the frequency aspect as sound intensity increases.

We hypothesized that temporal masking between syllable elements must give rise to distortions to speech and this plays a critical role in perceiving speech at various sound levels. By removing this temporal masking via our experimental paradigm, we demonstrated that indeed there were strong effects on performance, with control animals benefitting overall and HL animals benefitting through a mitigation of rollover. These results further lend support to the idea of the balance between audibility and distortions at high sound intensities and that hearing-impaired individuals could innately benefit from removing the distortions caused by temporal masking. One possible study could be performed in the future to test this such as creating natural speech sounds with ramps at the consonant-vowel junctions and observing if both normal-hearing and hearing-impaired listeners could benefit from this sound manipulation at high sound intensities. Further studies beyond the scope of this paper are needed to fully elucidate the range of effects in temporal masking of speech and whether this extends to a macroscale of full word or sentence recognition.

## Acknowledgements

This work was supported by a Wellcome Trust Senior Research Fellowship [200942/Z/16/Z] and a Human Frontier Science Program Fellowship [LT000036/2019]. We thank C.C. Lam for his assistance and advice.

